# Influence of fecal collection conditions and 16S rRNA gene sequencing protocols at two centers on human gut microbiota analysis

**DOI:** 10.1101/175877

**Authors:** Jocelyn Sietsma Penington, Megan A S Penno, Katrina M Ngui, Nadim J Ajami, Alexandra J Roth-Schulze, Stephen A Wilcox, Esther Bandala-Sanchez, John M Wentworth, Simon C Barry, Cheryl Y Brown, Jennifer J Couper, Joseph F Petrosino, Anthony T Papenfuss, Leonard C Harrison The ENDIA Study Group

**Affiliations:** The Walter and Eliza Hall Institute of Medical Research, Parkville, Victoria 3052, Australia; School of Medicine, University of Adelaide Robinson Research Institute, Adelaide 5006, South Australia, Australia; Alkek Center for Metagenomics and Microbiome Research, Department of Molecular Virology and Microbiology, Baylor College of Medicine, Houston, TX 77030-3411

**Keywords:** human, fecal, gut, microbiota, sample collection, 16S rRNA gene, QIIME

## Abstract

**Background:** To optimise fecal sampling and analysis yielding reproducible microbiome data, and gain further insight into sources of its variation, we compared different collection conditions and 16S rRNA gene sequencing protocols in two centers. Fecal samples were collected on three sequential days from six healthy adults and placed in commercial collection tubes (OMNIgeneGut OMR-200) at room temperature or in sterile 5 ml screw-top tubes in a home fridge or home freezer for 6-24 h, before transfer at 4°C to the laboratory and storage at - 80°C within 24 hours. Replicate samples were shipped on dry ice to centers in Australia and the USA for DNA extraction and sequencing of the V4 region of the 16S rRNA gene, using different PCR protocols. Sequences were analysed with the QIIME pipeline and Greengenes database at the Australian center and with an in-house pipeline and SILVA database at the USA center.

**Results:** Variation in gut microbiome composition and diversity was dominated by differences between individuals. Minor differences in the abundance of taxa were found between collection-processing methods and day of collection. Larger differences were evident between the two centers, including in the relative abundances of genus *Akkermansia*, in phylum *Verrucomicrobiales*, and *Bifidobacteria* in *Actinobacteria*.

**Conclusions:** Collection with storage and transport at 4°C within 24 h is adequate for 16S rRNA analysis of the gut microbiome. However, variation between sequencing centers suggests that cohort samples should be sequenced by the same method in one center. Differences in handling, shipping and methods of PCR gene amplification and sequence analysis in different centers introduce variation in ways that are not fully understood. These findings are particularly relevant as microbiome studies shift towards larger population-based and multicenter studies.

## Background

The fecal or ‘gut’ microbiome is shaped strongly by diet but also by the host genotype, age, hygiene and antibiotic exposure, and is altered in many pathophysiological states [1], [2]. The composition of gut microbiota differs greatly between individuals [3] and therefore maximizing the detection of biological and disease-related changes requires minimization of variation due to methods of sample collection and analysis.

Previous studies have shown that fecal microbial composition overall was not altered when DNA was extracted from a fresh fecal sample compared to a sample that had been immediately frozen and stored at −80 °C for up to 6 months [4, [5]. Storage at different temperatures for varying times has been compared with immediate freezing and storage at −80 °C. One study [5] reported a decrease in *Bacteroidetes* and an increase in *Firmicutes* phyla after 30 minutes at room temperature, but the majority [4, 6-11] have found that storage at room temperature for at least a day, or at 4, −20 or −80 °C for up to 14 days, had little effect on the relative abundance of taxa. Moreover, microbial composition was not significantly altered in DNA extracted from fecal occult blood test cards that had been at room temperature for three days [12]. Recent studies have also evaluated the OMNIgene®•GUT (0MR-200) collection and liquid storage tube, which is reported to stabilize DNA at room temperature for 14 days [13]. Samples immediately collected into these tubes and stored for three days at room temperature showed little difference in microbial composition by 16S rRNA gene sequencing compared with samples immediately frozen at minus 80 °C [7]. A similar result was obtained when samples in the tubes, stored for 1-28 days at room temperature, were compared with corresponding samples from which DNA had been freshly extracted [14, [15]. However, the relative abundance of *Bacteroides* increased after seven days and in infants (with lower microbial diversity than adults) significant divergence from fresh samples was observed after 14 days storage.

As microbiome studies expand into larger populations at multiple sites, stringent quality control remains critical. With this in mind and to optimise analysis of the gut microbiome in a multi-site, longitudinal pregnancy-birth cohort study [16], we evaluated different collection-processing methods on three sequential daily fecal samples from six individuals, and 16S rRNA gene sequencing in two centers.

## Methods

Six healthy adult volunteers, three males and three females aged 35-70, provided fecal samples on three successive days. Multiple aliquots were taken from each bulk sample and those for bacterial microbiome analysis were stored by one of four methods A-D (Figure 1).

**Figure 1.**
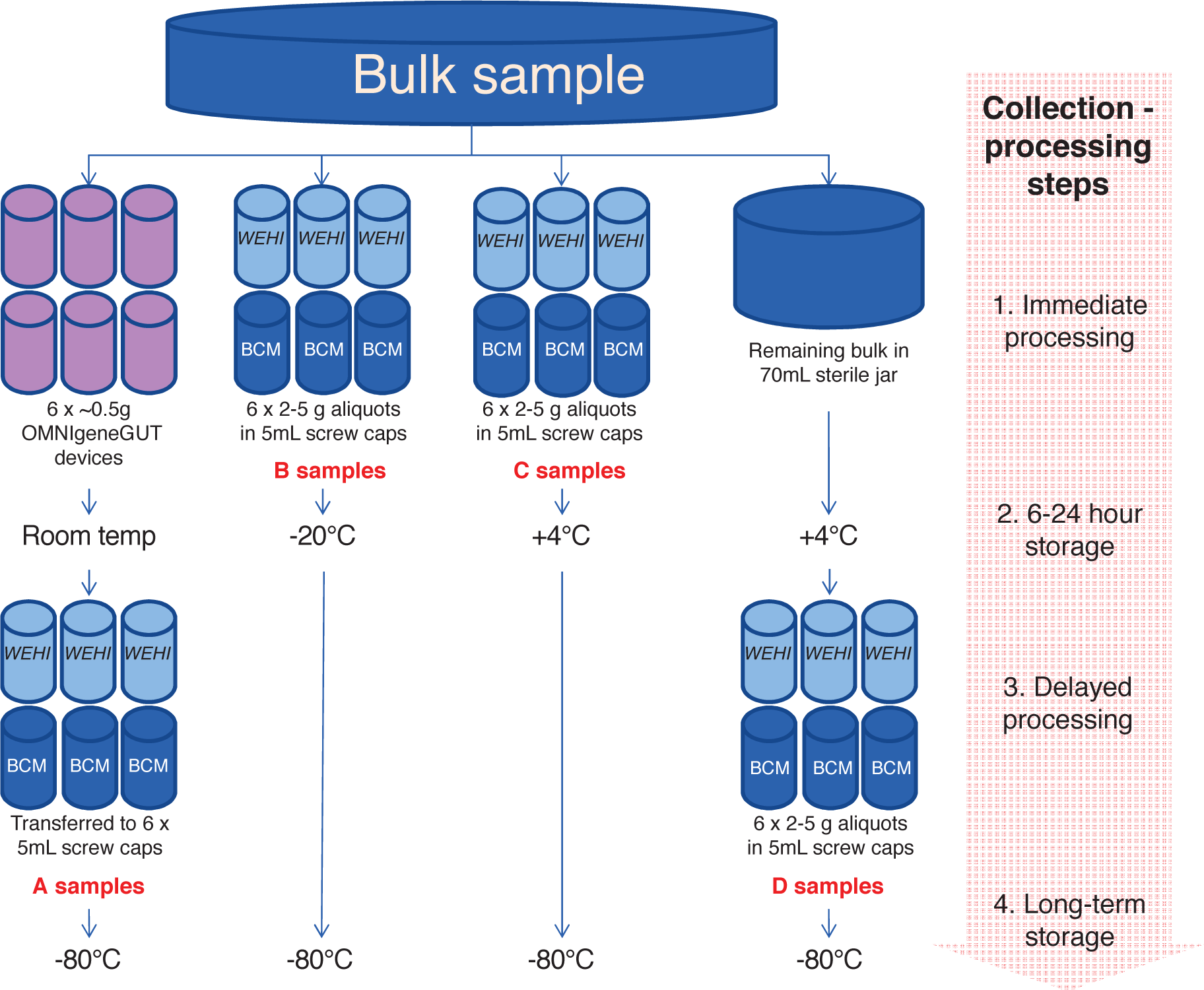
Schematic of the four collection methods for a home-collected fecal sample. Six individuals collected samples over three days. Each sequencing center received three aliquots per person-day-method combination (216 in total).

### Collection-processing methods

Method A: individuals placed aliquots of feces into 6 × OMNIgene®•GUT (0MR-200) [13] tubes as per manufacturer‘s protocol. These were stored at room temperature for 6-24 h before delivery to the laboratory for transfer to sterile 5 mL screw cap tubes and storage at minus 80°C. Method B: individuals placed aliquots of feces into 6 × sterile 5 ml screw cap tubes, which were stored in the home freezer for 6-24 h before delivery to the laboratory in an insulated container for storage at minus 80°C. Method C: individuals placed aliquots of feces into 6 × sterile 5 m screw cap tubes, which were stored in the home refrigerator for 6-24 h before delivery to the laboratory in an insulated container for storage at minus 80°C. Method D: individuals placed a bulk fecal sample into a sterile 70 ml collection jar, which was stored in home refrigerator for 6-24 h before delivery to the laboratory in an insulated container, transfer of aliquots into 6 × sterile 5 m screw cap tubes and storage at −80°C.

In the laboratory, samples were handled under sterile conditions in a Biosafety Level 2 cabinet. This collection-processing procedure yielded a total of 432 samples (24 from each of 6 individuals per day for 3 days) (Figure 1). After 2-4 weeks at −80°C, three sample aliquots per person-day-method (n=216) were transported on dry ice to sequencing centers in Australia (Walter and Eliza Hall Institute of Medical Research, Melbourne, Victoria; WEHI), and the USA (Baylor College of Medicine, Alkek Center for Metagenomics and Microbiome Research, Houston, Texas; BCM), for 16S rRNA gene amplicon sequencing. At both WEHI and BCM samples were stored at −80°C for 4 weeks before sequencing.

### DNA sequencing methods

Samples were thawed on ice and DNA extracted at both WEHI and BCM with the MoBio PowerSoil kit (MoBio Laboratories, Carlsbad, CA), as used in the Human Microbiome Project [17]. At WEHI, the V4 hypervariable region of the bacterial 16S rRNA marker gene (16Sv4) was PCR-amplified in duplicate with primers 515F-OH1 and 806R-0H2. Analogues of these are described, respectively, by the common name U515F, new name S-*-Univ-0515-a-S-19, and the common name 806R, new name S-D-Bact-0787-b-A-20 in [18]. These primers (<u>GTGACCTATGAACTCAGGAGTC</u>GGACTACNVGGGTWTCTAAT) and (<u>CTGAGACTTGCACATCGCAGC</u>GTGYCAGCMGCCGCGGTAA) included unique sequences (underlined) that provide a target for subsequently introducing Illumina sequencing adaptors and dual index barcodes to the amplicon target for paired-end sequencing on the Illumina MiSeq instrument [19]. Primary 16S rRNA gene amplification PCR cycling conditions were: 94°C for 3 minutes followed by 20 cycles at 94°C for 45 seconds each, 55°C for 1 minute, and 72°C for 1 minute 30 seconds and a final extension step at 72°C for 10 minutes. Successful amplification was determined by agarose gel electrophoresis. Amplicons from the primary amplification were diluted 1/10 and used as template for the secondary amplification. In the secondary amplification, the overhang sequences were used to introduce Illumina sequencing adaptors and dual index barcodes to the amplicon target. Individual amplicons were identified using 8 base index sequences from the Illumina Nextera design. Sixteen forward index primers and 24 reverse index primers were designed for a 96-well plate format with the potential to generate a maximum of 384 dual index amplicons (SRT1_OH1 – 5’-CAAGCAGAAGACGGCATACGAGATCCGGTCTCGGCATTCCTGCTGAACCGCTCTTCCGATCTNNNNNNNNGTGACCTATGAACTCAGGAGTC-3’ and SRT2-OH2 – 5’-AATGATACGGCGACCACCGAGATCTACACTCTTTCCCTACACGACGCTCTTCCGATCTNNNNNNNNCTGAGACTTGCACATCGCAGC-3’. The sequence NNNNNNNN indicates where individual indexes are placed in the oligo design). PCR conditions were as for the primary amplification, except for an increase to 25 cycles. Reactions were performed in triplicate in separate plates, with extra wells added if PCR product appeared low. Amplicon size distribution was determined by agarose gel electrophoresis. One sample failed to yield sufficient product in any well. Reactions from the three replicate plates were pooled and the PCR amplicons purified with 1.0× NGS Beads (Macherey-Nagel). Each dual indexed library plate pool was quantified with the Agilent Tapestation and the Qubit™ DNR BR assay kit for Qubit 3.0^®^ Fluorometer (Life Technologies).The indexed pool was diluted to 12pM and sequenced with the paired end 600-cycle (2×311) kit on a MiSeq instrument. After quality filtering, 702 technical replicate samples comprising 215 biological samples from the 6 individuals were obtained.

At BCM, the 16Sv4 region was also PCR-amplified with primers 515F and 806R and a singleend barcode sequence for each sample. 16S rRNA gene sequencing methods were adapted from the NIH-Human Microbiome Project [17] and the Earth Microbiome Project [20, [21]. The primers used for amplification contained adapters for MiSeq sequencing and single-end barcodes allowing pooling and direct sequencing of PCR products [21]. PCR-amplified amplicons were normalized by concentration before pooling and sequences were generated in one lane of a MiSeq instrument using the v2 kit (2×250 bp paired-end protocol). The 216 samples were analysed in a single pool.

### Bioinformatics methods

Two different bioinformatics pipelines were applied to the sequence data, Pipeline W at WEHI and Pipeline B at BCM. In Pipeline W, sequences were clustered into operational taxonomic units (OTUs) with 97% similarity using QIIME (Version 1.8.1) [22-25], and taxonomically classified by aligning the representative sequences to the Greengenes 13_08 database [26]. Paired-end sequences were assembled with PEAR [27] with parameters -v 100 -m 600 -n 80, where -v is minimum overlap, -m is maximum assembled length and -n is minimum assembled length. WEHI index sequences were extracted with QIIME script extract_barcodes.py, and bases up to and including the 533F to 805R V4 region amplicon primers were trimmed. BCM sequences were supplied as trimmed sequences with a separate barcode index file.

QIIME’s split_libraries_fastq.py script was used for quality filtering, with phred quality scores required to be above 29, and 90% of a read’s length required to have consecutive, high-quality base calls. In order to minimise differences between WEHI and BCM sequences, the WEHI sequences were further filtered by aligning to the SILVA 123 16S rRNA gene database using MOTHUR v1.38.1 [28, [29] with start=8390 and end=17068, and removing sequences that were not in this position.

QIIME’s open-reference OTU-picking was used with the Uclust algorithm [22, [23] to form clusters at 97% similarity. The representative sequences from each OTU were aligned with gaps to a reference set using QIIME’s implementation of PyNAST, then filtered for chimeric sequences using UChime with default settings [30].

After making filtered OTU table with minimum count 3, and assigning taxonomy with Greengenes_13_08 [26], the data were analyzed in R.

WEHI data consisted of 759 (PCR-well) samples. Those with sequence counts < 1000 or with descriptions of PCR success of ‘None’ or ‘Very faint’ were discarded, which included all samples for individual 55, method B, day 3, aliquot 3. OTU counts for the remaining technical replicates were summed to give 215 biological-replicate samples.

Pipeline W was applied to sequence data from both WEHI and BCM centers. Results from Pipeline W applied to WEHI sequences are termed WW; results from Pipeline W applied to BCM sequences are termed WB. Alpha and beta diversity and differential abundance analyses were performed in R using the Phyloseq package [31] and DESeq2 [32], Most analysis used 215 or 216 samples with minimum OTU size of 20; alpha diversities were calculated with the minimum-count=3 OTUs with technical and biological replicates combined, giving 72 samples.

In Pipeline B, read pairs were de-multiplexed based on the unique molecular barcodes using Cassava 1.8.4 from Illumina, and reads were merged using USEARCH v7.0.1090 [23], allowing zero mismatches and a minimum overlap of 50 bases. Merged reads were also trimmed at first base with Q5. In addition, USEARCH was also used to apply a quality filter to the resulting merged reads and reads containing above 0.05 expected errors were discarded.

16S rRNA gene sequences were clustered into OTUs at a similarity cutoff value of 97% using the UPARSE algorithm [33]. OTUs were mapped to an optimized version of the SILVA Database [18] containing only the 16Sv4 region to determine taxonomies. A rarefied OTU table was constructed from the output files generated in the previous two steps for downstream analyses of alpha diversity, beta diversity and phylogenetic trends [34]. Pipeline B was applied only to BCM data.

## Results

For WW protocol, library sizes after quality filtering, clustering, and combining PCR replicates ranged from 30,000 to 250,000 sequences per sample, with a median of 67,000 (Figure S1A); sequences clustered into 12,652 OTUs of minimum size 20. For WB, library sizes ranged from 5,000 to 56,000, with a median of 27,000 (Figure S1B); sequences clustered into 3,675 OTUs of minimum size 20. The differences between library sizes reflect the differences between technical replicates (three for WEHI, one for BCM), as well as sequencing and filtering differences.

### Taxonomic overview

At the phylum level, samples were dominated, as expected, by *Bacteroidetes* and *Firmicutes*. The mean summed proportion of these two phyla was 94%, varying from 71.9% to 99.7% between individual samples. Likewise, a single order from each of these two phyla, *Bacteroidales* and *Clostridiales*, was dominant, with three orders from the phylum *Proteobacteria* contributing another 1-2% overall (Figure 2A; see also Figure S2A).

**Figure 2.**
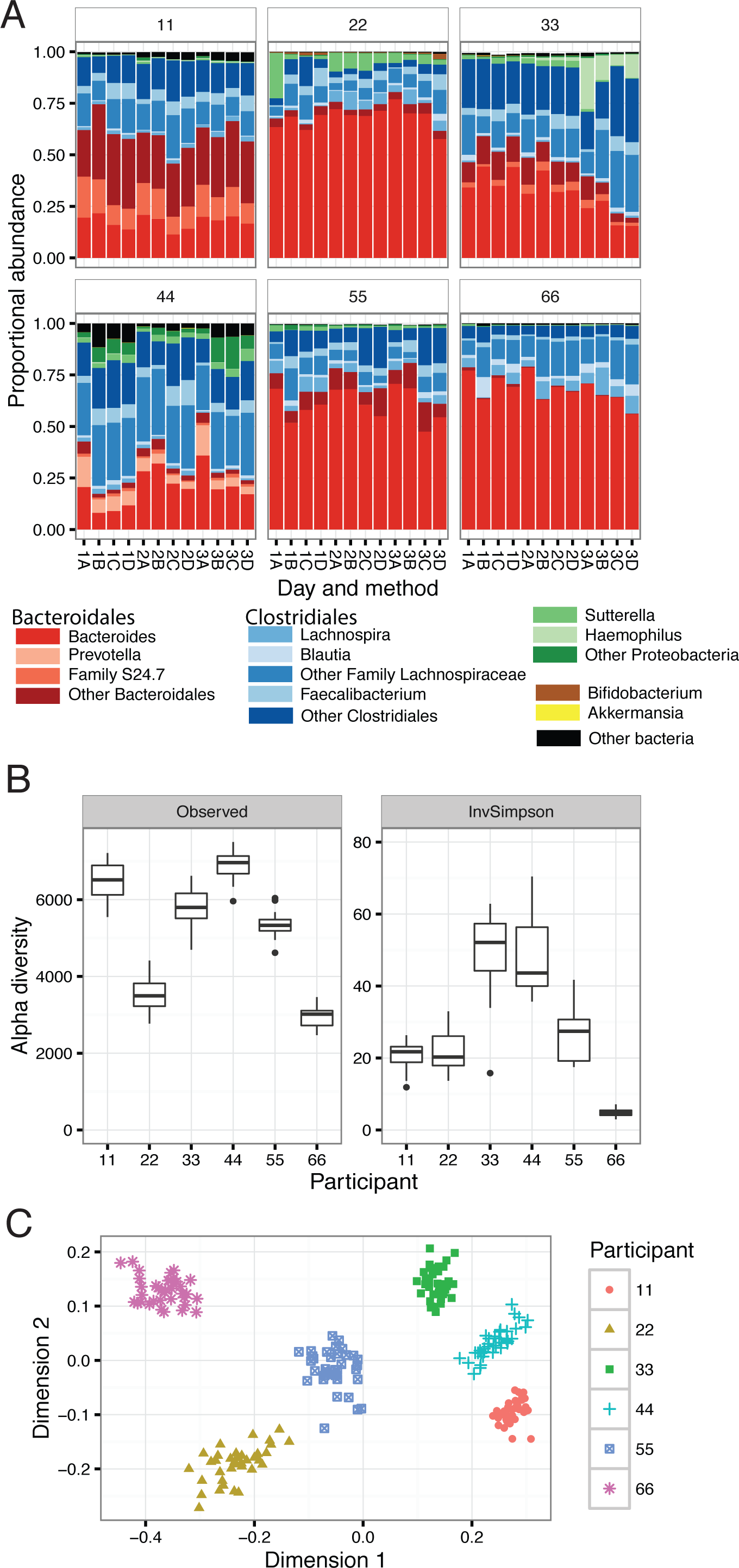
Overview of the fecal bacterial microbiome, WW protocol. (A) Dominant bacterial genera in fecal samples, or higher taxa where genus was not available. Bars are colour-coded by phyla: red *Bacteroidetes*, blue *Firmicutes*, green *Proteobacteria*, brown *Actinobacteria*, yellow *Verrucomicrobia*. (B) Alpha diversity within samples. Two measures are shown: observed number of OTUs per sample, an estimate of richness, and Inverse-Simpson index indicating the evenness of the sample. Samples were sub-sampled to the smallest sample size of 108,000 sequences, and values are the mean of 10 random sub-samples. Boxes show the interquartile range for the four methods on three days. (C) Beta diversity. NMDS ordination of the UniFrac distance between samples, a representation of phylogenetic similarity.

Alpha (α) diversity is used to characterise the richness of the microbiome and its evenness (heterogeneity) or distribution of proportions. Samples showed a considerable spread of α diversity (Figure 2B). Samples from individual 66 had the lowest observed richness (number of OTUs per sample) and the lowest Inverse-Simpson diversity index, the latter indicating dominance by a smaller number of OTUs. This is reflected in the genera plots (Figure 2A). In contrast samples from individual 11 had a high observed richness but a comparatively low Inverse-Simpson index, consistent with the presence of a few high-abundance and multiple low-abundance genera.

Analysis of β diversity by non-metric multidimensional scaling (NMDS) ordination of the UniFrac distance showed that samples cluster strongly by individual, with marked separation between individuals (Figure 2C).

### Effect of collection-processing method on taxonomic analysis

With the WW protocol, testing for differential abundances between collection-processing methods revealed no significant differences in counts by phylum, order or family (Table 1, Figure 3A). The differences were tested using DESeq2 with a design controlling for the effect of person and day. A very small number of OTUs (0.04%) were different under collection-processing Method A.

**Figure 3.**
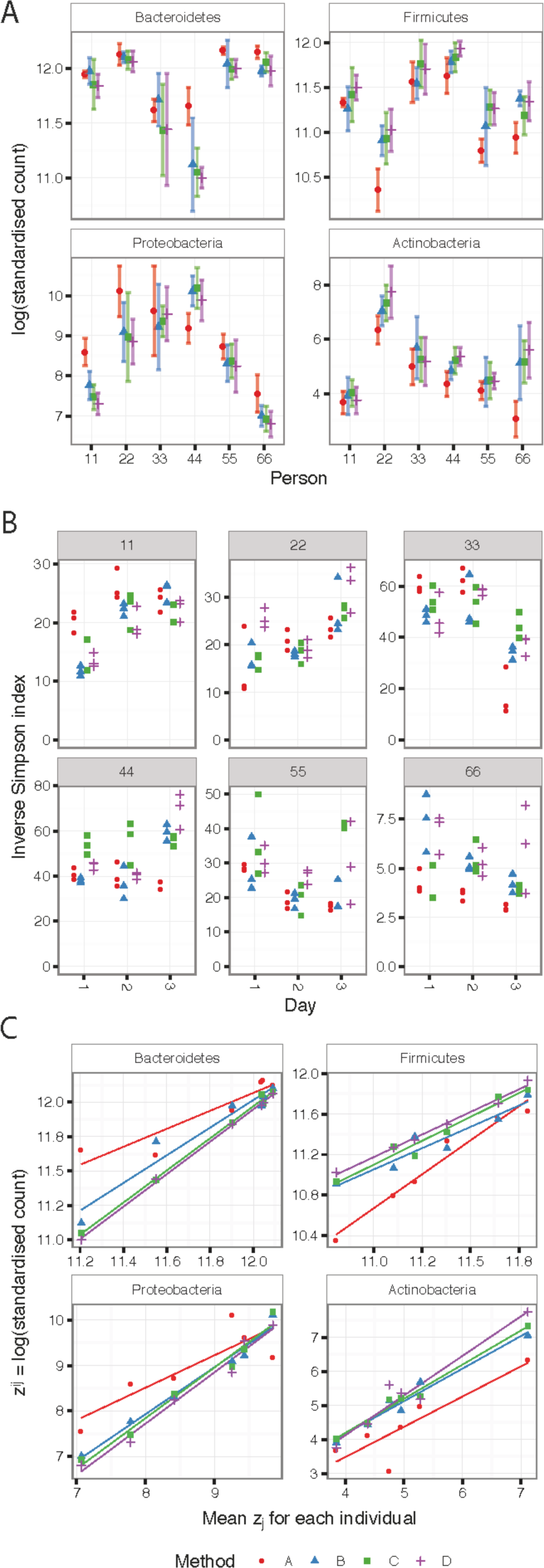
Effect of collection-processing method by analysis WW. (A) Log of standardised counts (scaled by library size) of the four most abundant phyla. Points show mean and bars standard deviation (sd) for each individual and collection-processing method. Method A has the smallest average sd for *Bacteroidetes* and *Actinobacteria*. (B) The Inverse-Simpson α diversity index for each sample (compare with Fig 2B). (C) Mean log (standardised count) plotted against the mean over the collection-processing methods, and a linear regression applied. Method A has the greatest average deviation from the linear model.

**Table 1.**
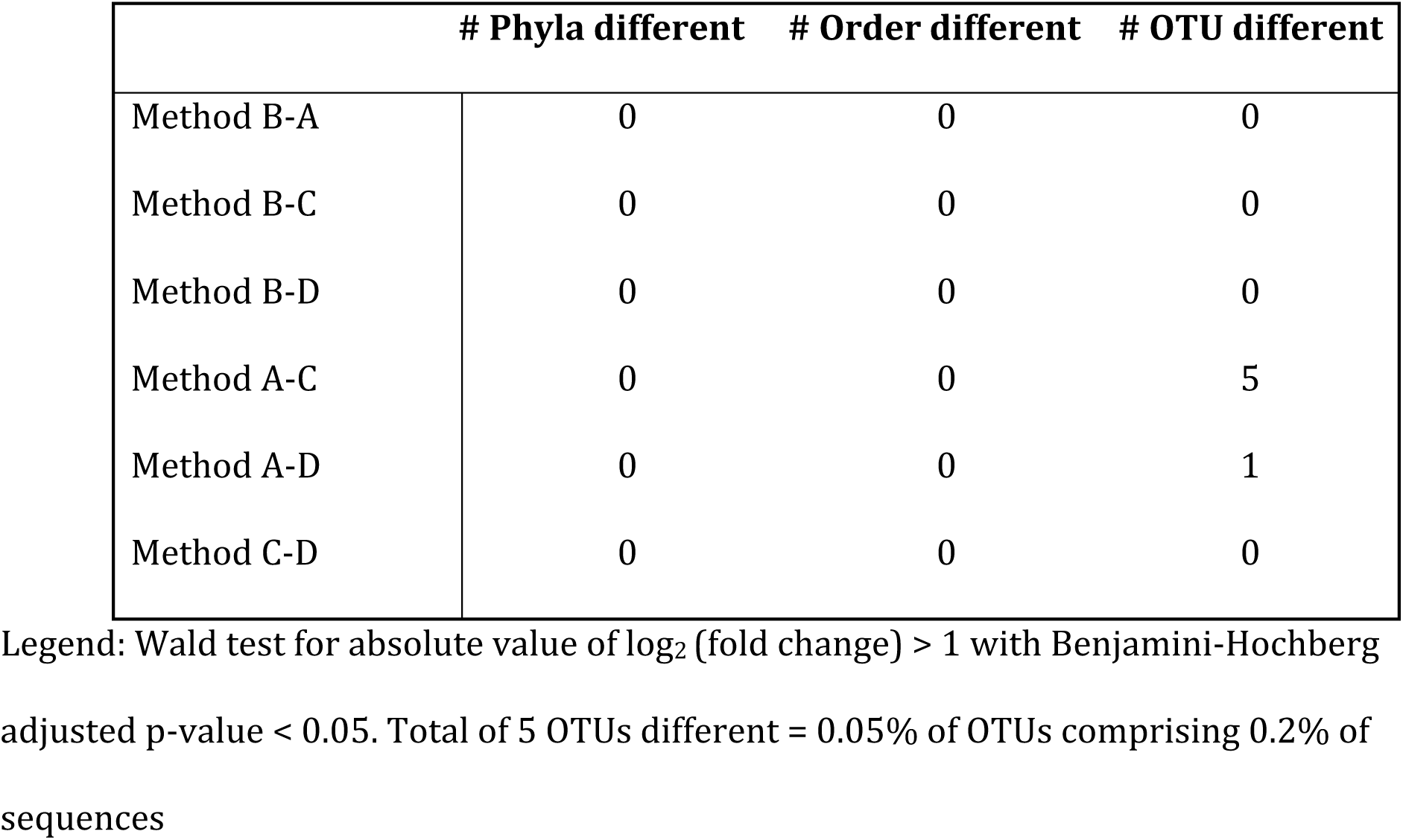
Differences by collection-processing method at three taxonomic levels (analysis WW).

Diversity varied within a sample depending on collection-processing method (Figure 3B) but the effect was small and inconsistent. After fitting a linear model with inputs for method and individual, 23% of variation was unaccounted for while collection-processing method accounted for only 2%. Overall, alpha diversity was slightly lower from Method A (Table 2).

**Table 2.** Change in α diversity (Inverse-Simpson index) due to collection-processing method (WW protocol).

| **Method comparison** | **95% confidence lower bound** | **95% confidence upper bound** | **Family-wise adjusted p value** |
| --- | --- | --- | --- |
| **A-B** | -4.9 | 3.2 | 0.95 |
| **C-B** | -0.5 | 7.6 | 0.10 |
| **D-B** | -1.1 | 6.9 | 0.24 |
| **C-A** | 0.4 | 8.4 | 0.02 * |
| **D-A** | -0.2 | 7.8 | 0.07. |
| **D-C** | -4.6 | 3.4 | 0.98 |

Different methods of collection-processing might also increase the variance between samples, reducing the reproducibility of a result. Two approaches were used to test for this. Greater variance between samples is equivalent to greater distance between samples by some measure. The Bray-Curtis dissimilarity between OTU counts was calculated for pairs of samples from each individual and method, and a Tukey Honest Significant Difference test applied to a linear model of the dissimilarity. There was no evidence that the dissimilarity between samples was different for collection-processing methods (smallest p=0.1). In addition, we looked for differences in the variance of the four most abundant phyla. The log transformed standardised counts for *Bacteroidetes*, *Firmicutes*, *Proteobacteria* and *Actinobacteria* per sample were compared with the mean across collection-processing methods for each individual (Figure 3C). Methods B, C and D gave similar results, while method A had lower variance within samples from the same individual but greater deviation from the mean compared with the other methods.

### Effect of collection-processing method on library size

Collection-processing methods were compared after quality filtering, barcode extraction and clustering. Methods did not differ significantly in the number of sequences identified by the WW protocol. Batch effects were more significant (p<10^-5^) than collection-processing method, but batch and method together contributed less than 5% of the variation in library size. In summary, different collection-processing methods for fecal samples were associated with only minor differences in the microbiota predicted by 16S rRNA gene sequencing. Method A with the OMNIgeneGut OMR-200 device had minor taxonomic differences compared with Methods B-D that involved immediate freezing or refrigeration. The number of DNA sequences extracted per sample was not different by collection-processing method, and no major differences in abundance were apparent at the phylum level. In the WW analysis, no method was associated with significantly more distance between samples at the OTU level. Thus, different collection-processing methods were associated with minor variations but no consistent bias. Overall, the differences between individuals were much larger than those introduced by the different collection and processing methods.

### Effect of sequencing center

Sequences generated at WEHI and BCM were analyzed with pipeline W. The most abundant phyla were similar, but the proportions of less abundant phyla and higher taxonomic resolution differed. For example, the mean proportion of genus *Akkermansia* in the order *Verrucomicrobiales* was greater in WB (0.7%) than WW (0.02%). The proportion of *Bacteroides* was lower in some individual samples for BCM than WEHI (Figure 4A, also S2).

**Figure 4.**
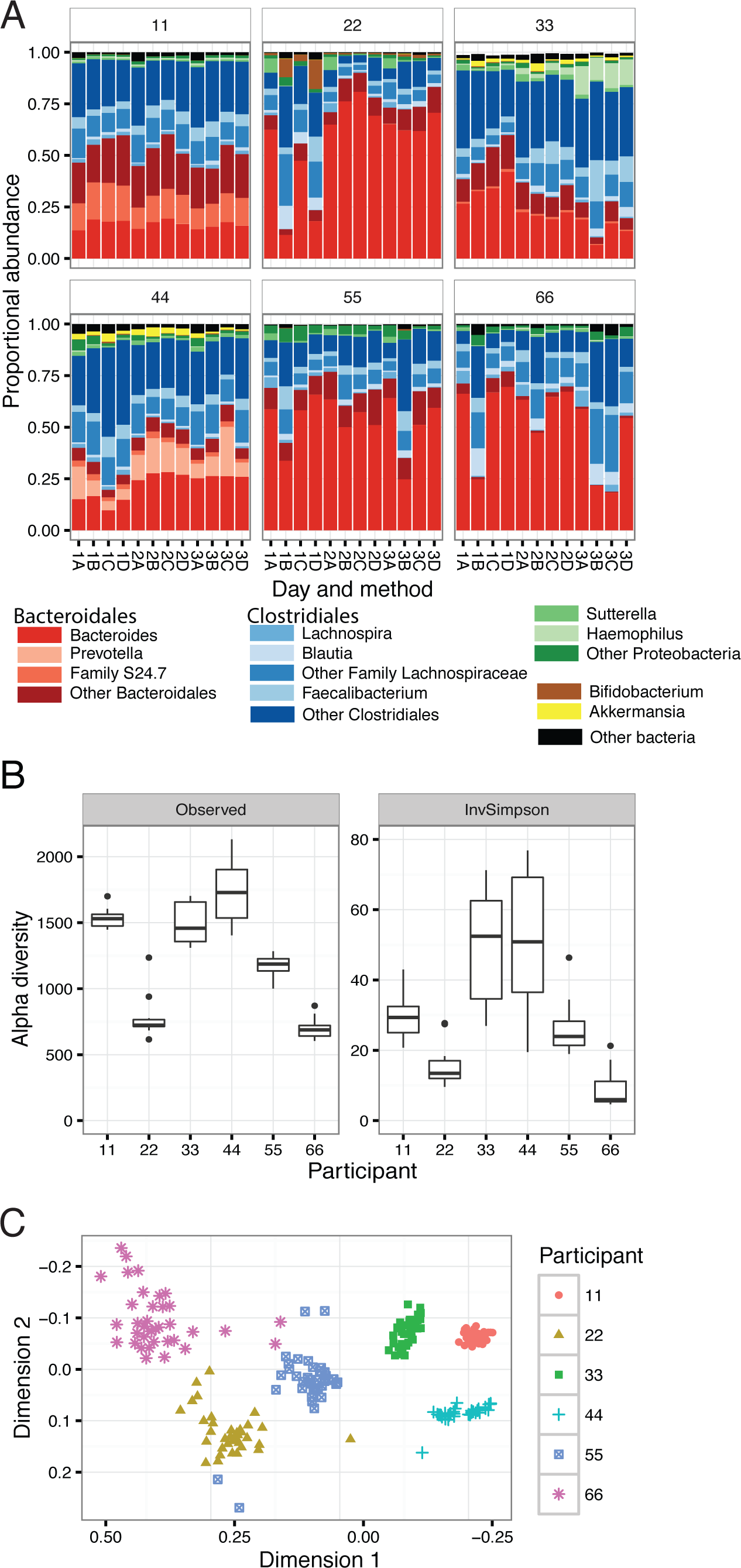
Overview of the fecal bacterial microbiome, WB protocol. (A) Dominant bacterial genera in each sample. Bars are colour-coded by phyla as in Figure 2. (B) Alpha diversity within samples, sub-sampled to 31,500 sequences. (C) Beta diversity as NMDS ordination of UniFrac distance.

WB contained fewer OTUs and therefore lower values for observed richness (Figure 4B). The number of OTUs observed per sample was dependent on sampling depth (Figure S3); values shown are based on the smallest sample sizes for each of the two data sets. Richness was similar between the WEHI and BCM data sets analysed by pipeline W, with samples from individual 66 showing the lowest alpha diversity and those from individual 44 the highest (Figure 2B, 4B). For the Inverse-Simpson diversity index, which is not dependent on library size at this depth of sequencing, WB had a greater range of values, and a greater range for samples from some individuals. WW and WB had similar patterns of beta diversity between individuals (Figures 2C and 4C), although WB had several outliers.

Initial bioinformatics analysis was performed separately on the WEHI and BCM datasets. For better comparison of the taxonomies, pipeline W was re-applied to a data set comprising the BCM sequences and one of the three WEHI technical replicates (Figure 5). The ordination plot shows ‘batch’ effects between the two sequencing centers, and greater between-sample differences in the BCM data.

**Figure 5.**
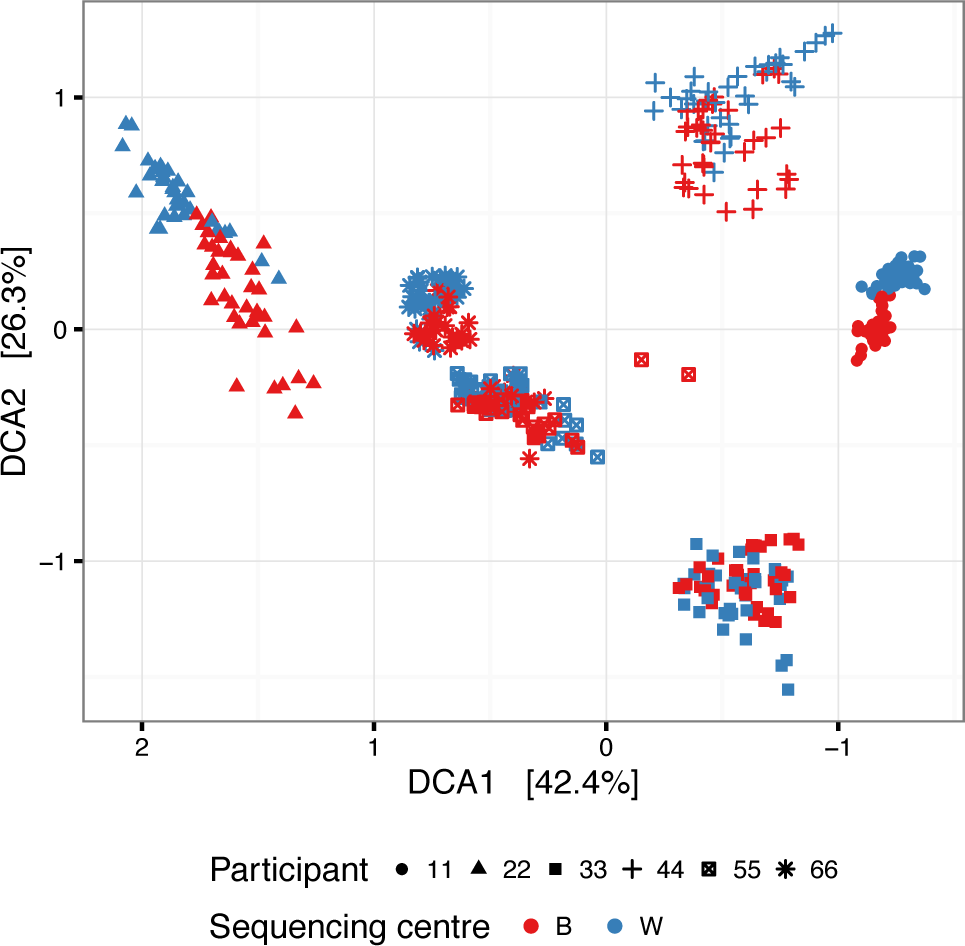
Beta diversity between samples from two sequencing centers. Ordination plot of Bray-Curtis distances between samples, using Detrended Correspondence Analysis. Points represent samples from WB and a single technical replicate from WW.

DESeq2 was used to make generalized linear models for the counts at phylum, order and OTU levels (Table 3). The model included individual ID, day and collection-processing method as factors. At the phylum level, the largest change was in the *Verrucomicrobia*. At the OTU level, 3% of OTUs were significantly different (Figure S4, Additional data S1). Most of the differentially abundant OTUs belonged to the orders *Clostridiales* (63%) and *Bacteroidales* (31%). The direction of change in OTUs was not consistent, and there were no significant differences in counts for *Clostridiales* and *Bacteroidales* between WEHI and BCM.

**Table 3.** Sequencing center comparisons: pipeline W applied to combined sequence sets.

| **Phylum** | Base Mean | Log<sub>2</sub> W/B | Adjusted p-value | Max proportion W | Max proportion B |
| --- | --- | --- | --- | --- | --- |
| *Verrucomicrobia* | 39.2 | -2.9 | 2.6E-39 | 1.1% | 5.1% |
| *Actinobacteria* | 68.7 | -1.5 | 3.5E-06 | 3.6% | 21.3% |
| *Lentisphaerae* | 9.6 | 1.6 | 1.5E-02 | 1.3% | 0.3% |
| **Order** |  |  |  |  |  |
| *Verrucomicrobiales* (V) | 38.7 | -3.2 | 4.8E-45 | 0.3% | 5.1% |
| *Enterobacteriales* (P) | 146.4 | -2.0 | 4.3E-10 | 8.4% | 9.7% |
| *Actinomycetales* (A) | 0.8 | -2.0 | 2.4E-07 | 0.03% | 0.1% |
| *Victivallales* (L) | 10.4 | 2.0 | 9.6E-06 | 1.3% | 0.3% |
| *Bifidobacteriales* (A) | 44.1 | -1.4 | 6.8E-03 | 3.5% | 20.9% |

**Legend:** Log<sub>2</sub> (fold change) is based on a fitted mean of the counts in samples as calculated by the DESeq2 package [32]. W: DNA sequenced at WEHI; B: DNA sequenced at BCM. The maximum proportions are included for context. The adjusted p-value is from a Wald test with the Benjamini & Hochberg false discovery rate. For each order the phylum is indicated in brackets: A *Actinobacteria*, L *Lentisphaerae*, P *Proteobacteria*, V *Verrucomicrobia*

With WB, collection-processing Methods A and B were taxonomically different, with a decrease in *Actinobacteria* in method A and an increase in *Lentisphaerae* (although counts were very low) (p < 0.001, Additional Table S1). *Lentisphaerae* were also increased in method A compared with methods C and D (p < 0.05). There were significant differences between Bray-Curtis distances between samples in WB data (p < 0.001), with collection-processing method A associated with smaller differences between samples from the same individual than methods B, C and D (Additional Table S2). Collection-processing method D resulted in fewer sequences than A or C (p=0.01) with the WB protocol, but the difference was small compared with total variation. (Figure S5).

### Effect of bioinformatics pipeline

BCM sequence data were also analysed with Pipeline B. This used the same fastq files of sequenced DNA of the 16Sv4 region as in WB, but allocated OTUs and assigned taxonomies according to a BCM protocol incorporating a version of the SILVA database [18]. Library sizes varied from 2,300 to 23,000 sequences per sample with a median of 14,000. Sequences were clustered into 463 OTUs, fewer than generated by Pipeline W. Pipeline W used the UCLUST algorithm for clustering, which is known to produce more OTUs [35]. The differences in OTU formation and classification resulted in similar descriptions to protocol WB at the phylum level, but with some differences at the order level (Figure S6A). The UniFrac distances clustered the samples in a very similar manner to protocol WB (Figure S6B). The effect of the collection-processing method was stronger than for WW (Figure S6C), and was similar to WB (Additional Table S3). Differential abundance analysis with DESeq2 indicated a difference in *Actinobacteria* counts between collection-processing methods, with method A giving lower proportions of *Actinobacteria* than B or C (p < 0.01). The different bioinformatic approaches gave very different OTU numbers, and some differences in microbiota designations due to differences between the reference databases, for example regarding the status of *Akkermansia*, and the polyphyletic *Clostridia*.

## Discussion

Collection-processing method had no discernable effect on microbiota taxonomy, or on the final number of 16S rRNA sequences counted, under protocol WW. There was a small but inconsistent effect on alpha diversity, with samples collected in the OMR-200 tubes (method A) having slightly lower average Inverse-Simpson indexes of diversity. The log-values of proportions of the four dominant phyla had lower variance within samples from the same individual, but greater deviation from the mean, for samples collected with method A.DNA extraction and sequencing at two centers, WEHI and BCM, showed some differences, particularly increases in the proportion of *Verrucomicrobia*, *Actinobacteria* and *Enterobacteriales* under protocol WB, compared with WW.

Under protocol WB, some effects of collection-processing methods were apparent. *Actinobacteria* proportions were reduced in Method A samples compared with the other collection-storage methods, and beta-diversity was also reduced. The reduction in distances between samples for an individual with collection-processing method A possibly indicates that the OMR-200 device mitigated the effects of storage and transport.

The differences between sequencing centers could be due to one or more of several factors. Although the samples were transported and arrived on dry ice, covert effects of shipping conditions can’t be excluded. Similarly, differences in sample handling at the laboratory level can’t be excluded. One clear difference between the centers was in the sequencing methods. Although both centers used the same primers for the primary PCR of the V4 amplicon of the 16S rRNA gene, the approaches then diverged. BCM barcoded the reverse primer for each sample and ran a one-step PCR with 32 cycles of amplification, whereas WEHI added overhang sequences to both primers to subsequently introduce both Illumina sequencing adaptors and dual index barcodes for each amplicon, and ran a two-step PCR with 45-cycles of amplification. In addition, BCM sequenced a single library pool whereas WEHI sequenced triplicated libraries in separate batches on three different days. These differences in methodology are likely to have contributed to inter-center differences in taxonomic composition.

## Conclusions

Collection-processing methods and day of collection contributed to only minor variation in fecal microbiome composition and diversity, the major variation being at the level of the individual. However variation, including at the phylum level, was evident between the two sequencing centers and is likely to be related at least in part to difference in PCR design and conditions. Collection with storage and transport at 4°C within 24 h is adequate for analysis of the gut microbiome, but variation between sequencing centers indicates that cohort samples should be sequenced by the same methods in one center. These findings are relevant to the quality control of microbiome studies, in particular to larger, population-based multi-site studies.

## Acknowledgements

We acknowledge Tulin Ayvaz, Ginger Metcalf, Donna Muzny and Richard Gibbs and the Human Genome Sequencing Center at Baylor College of Medicine (BCM) for assistance with sample processing and data generation for the BCM dataset.

## Declarations

### Ethics approval and consent to participate

Volunteers gave informed consent for self-collection of non-identifiable stool samples and basic, demographic information. In accordance with the Australian NHMRC National Statement on Ethical Conduct in Human Research, the research was considered as ‘negligible risk’ with no foreseeable risk of harm or discomfort to the volunteers. Accordingly, the investigation was exempt from review by the Human Research Ethics Committee.

### Availability of data and materials

Additional Figures S1 - S6 (pdf), Additional Tables Supp_Table1 - Supp_Table3 (docx) and Supp_data1, Supp_data2 (xlsx) attached.

R code is available as joint_plots_tables.Rmd from github.com/PapenfussLab/endiaQC/ Sequences are available through

https://www.ncbi.nlm.nih.gov/bioproject/?term=PRJNA393083

OTU tables are available through

https://figshare.com/articles/ENDIA_microbiome_QC_OTUs_and_metadata/5401594

## Author Contributions

JJC, LCH and MASP conceived the study. MASP, LCH, EB-S, JMW, SCB and CYB collected and processed samples. KMN, SAW and NJA performed DNA extraction and sequencing. JSP, AJR-S, ATP, NJA and JFP performed the bioinformatics analyses. JSP and LCH wrote the initial draft of the paper, which was reviewed by all authors.

## ENDIA Study Group

Peter G Colman^1^, Andrew Cotterill^2^, Maria E Craig^3,4^, Elizabeth A Davis^5,6^, Mark Harris^2^, Aveni Haynes^5,6^, Lynne Giles^7,8^, Grant Morahan^9^, Claire Morbey^10^, William D Rawlinson^4^, Richard O Sinnott^11^, Georgia Soldatos^12^, Rebecca L Thomson^7,13^, Peter J Vuillermin^14^

1 Department of Diabetes and Endocrinology, Royal Melbourne Hospital, Parkville, Victoria, Australia; 2 Endocrinology Department, Lady Cilento Children’s Hospital, Brisbane, Queensland, Australia; 3 Institute of Endocrinology and Diabetes, The Children’s Hospital at Westmead, Westmead, New South Wales, Australia; 4 Serology and Virology Division, NSW Health Pathology, Prince of Wales Hospital, Randwick, New South Wales, Australia, and University of NSW, Sydney, Australia; 5 Department of Endocrinology & Diabetes, Princess Margaret Hospital for Children, Perth, Western Australia, Australia; 6 Telethon Kids Institute, Subiaco, Western Australia, Australia; 7 Robinson Research Institute, University of Adelaide, Adelaide, South Australia, Australia; 8 School of Public Health, University of Adelaide, Adelaide, South Australia, Australia; 9 Centre for Diabetes Research, Harry Perkins Institute of Medical Research, Nedlands, Western Australia, Australia; 10 Hunter Diabetes Centre, Mereweather, New South Wales, Australia; 11 Melbourne eResearch Group, University of Melbourne, Parkville, Victoria, Australia; 12 Diabetes and Vascular Medicine Unit, Monash Health, Clayton, Victoria, Australia; 13 Adelaide Medical School, University of Adelaide, Adelaide, South Australia, Australia; 14 Child Health Research Unit, Barwon Health, Geelong, Victoria, Australia

---

**Figure S1.**
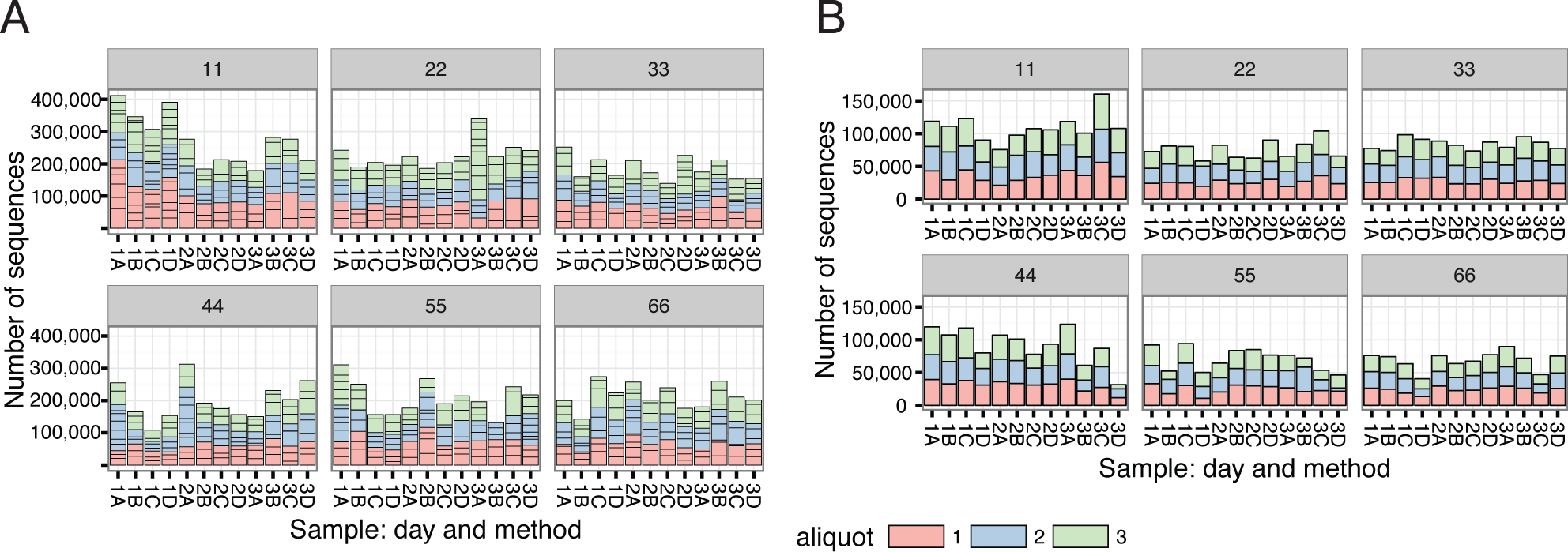
Number of sequences in combined samples. Colours indicate the three aliquots of each sample. (A) Walter and Eliza Hall Institute (WEHI) pipeline for WEHI 16S rRNA gene sequences (WW). Horizontal black lines within aliquots indicate the PCR replicates. Total number of sequences 15 million. (B) WEHI pipeline for Baylor College of Medicine Human Genome Sequencing Centre (BCM) 16S rRNA gene sequences. (WB). Total number of sequences 6.0 million

**Figure S2.**
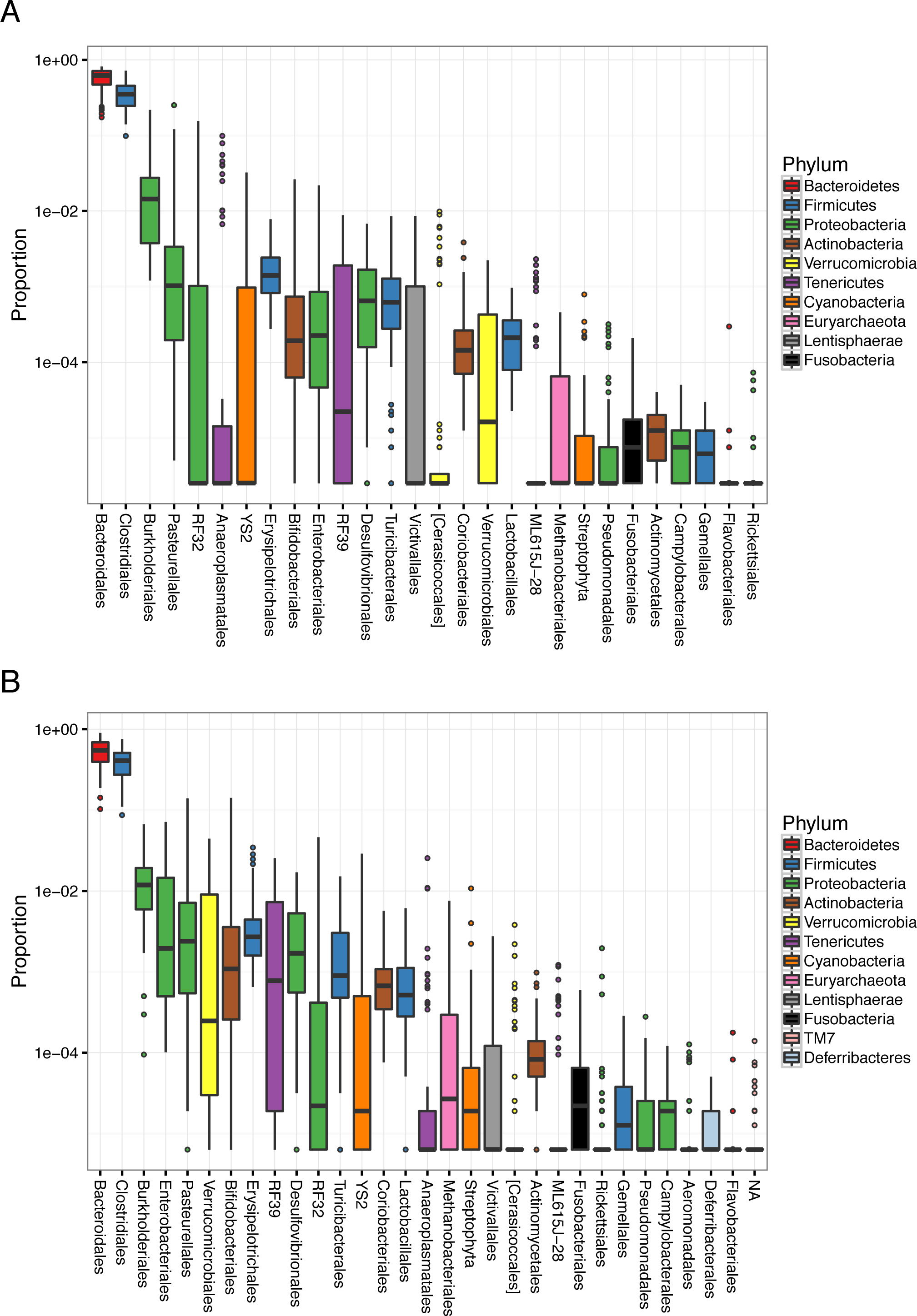
Proportion (log base 10) of bacterial orders in 72 samples (biological replicates combined). The X-axis is arranged by mean proportion. Order names are from Greengenes 13_08 database; square brackets indicate name proposed by the curators for an uncultured bacterium. Streptophyta is a chloroplast probably from undigested plant matter. (A) WW. (B) WB.

**Figure S3.**
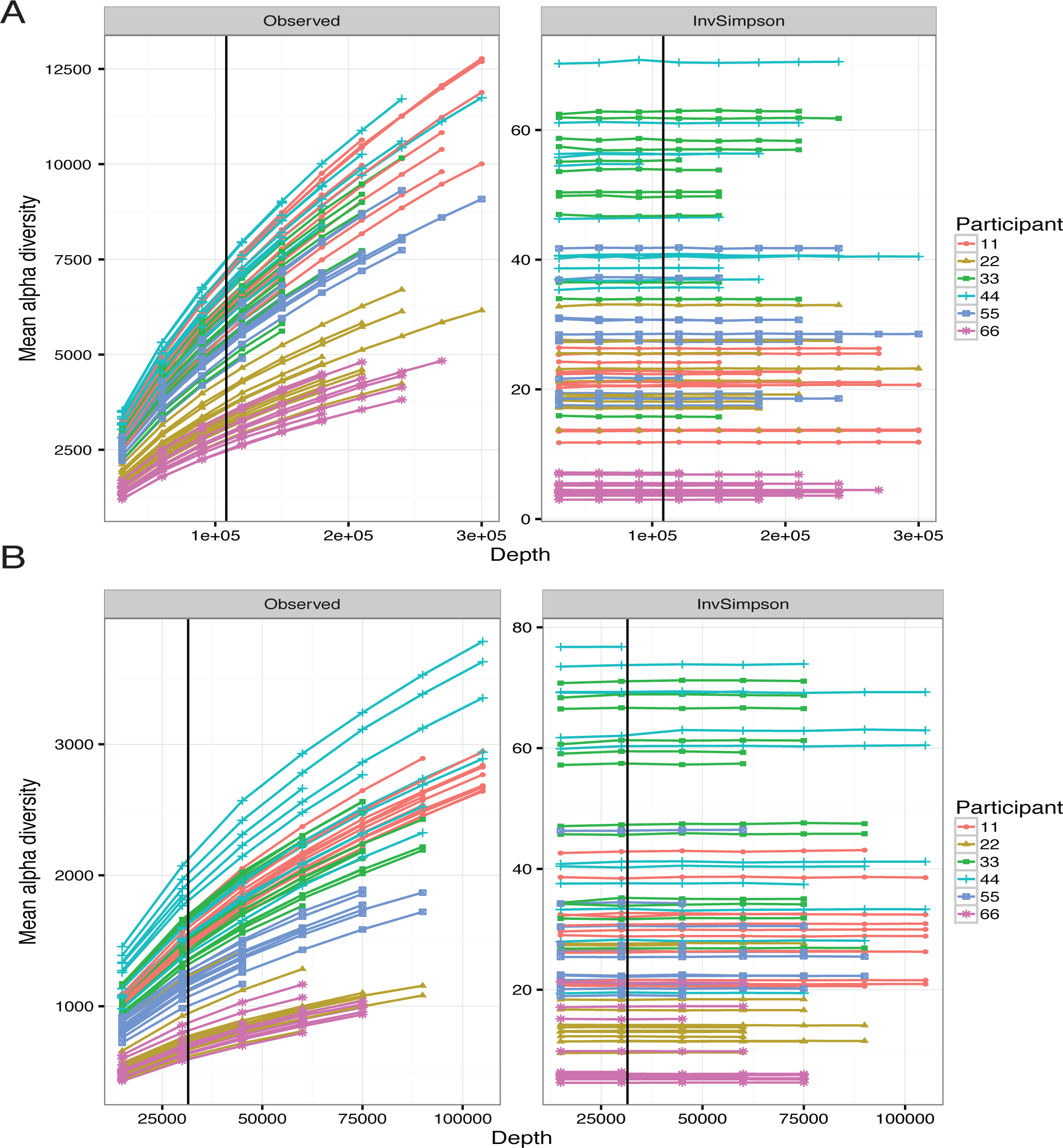
Two measures of sample α diversity at multiple sub-sampling depths for two sequencing facilities. Each individual has 12 plots, corresponding to four methods by three collection days. Values are means of 8 random sub-samples, taken without replacement. Standard deviation bars are smaller than point sizes. (A) WW data. The vertical black line shows the smallest sample. The smallest sample from WB is near the smallest sub-sample at 31,500 sequences. (b) WB data. The vertical black line shows the smallest sample.

**Figure S4.**
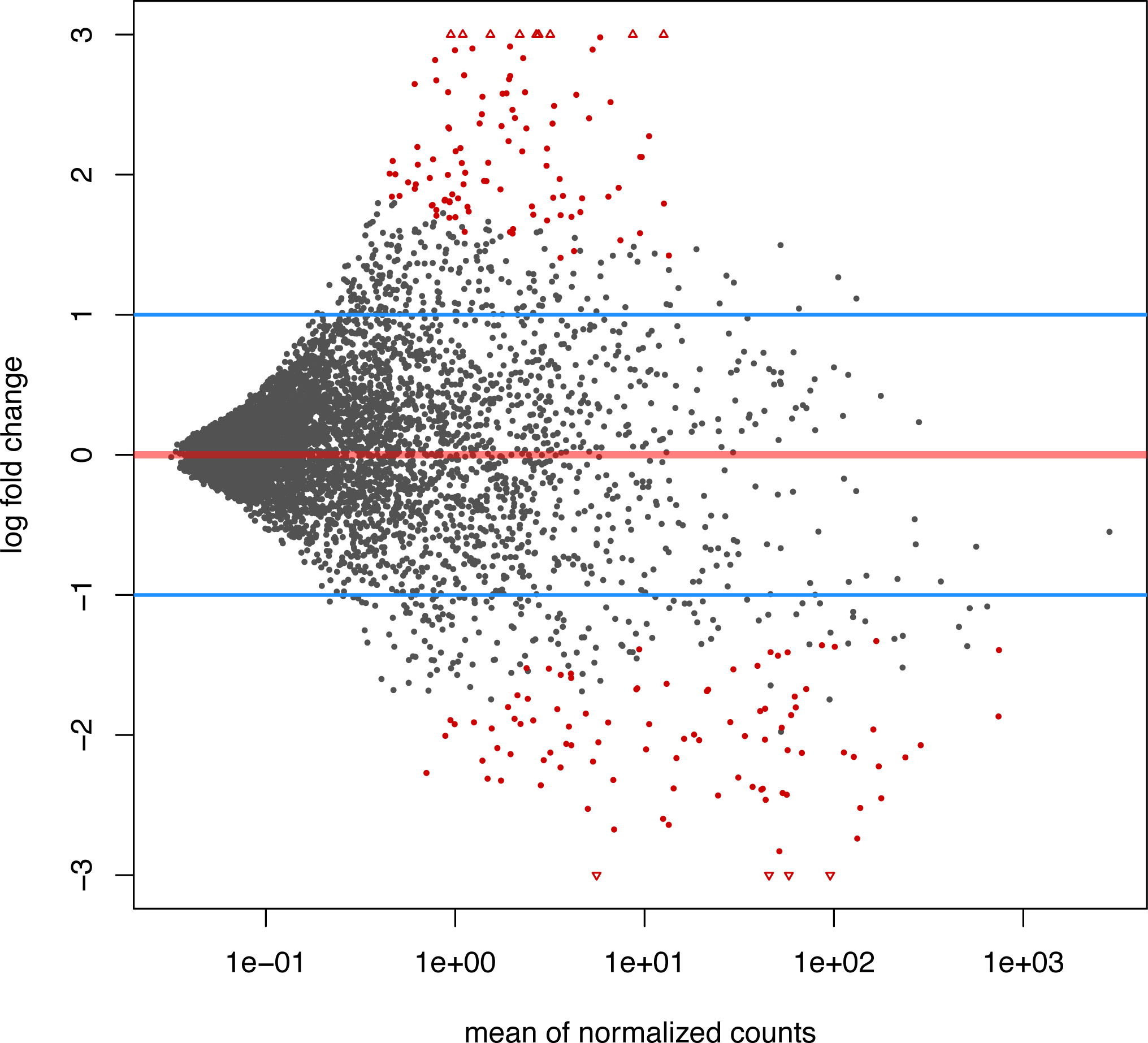
Scatter plot of OTUs showing log_2_ fold changes of W counts over B counts versus the mean normalized count. Red dots are OTUs with adjusted p < 0.05, triangles are points with fold changes outside the y-axis limits. The blue lines illustrate the null hypothesis that any change is less than 2-fold. Although the numbers of significant changes are evenly split between increase and decrease, W counts tend to be increased in small OTUs and B counts in larger OTUs.

**Figure S5.**
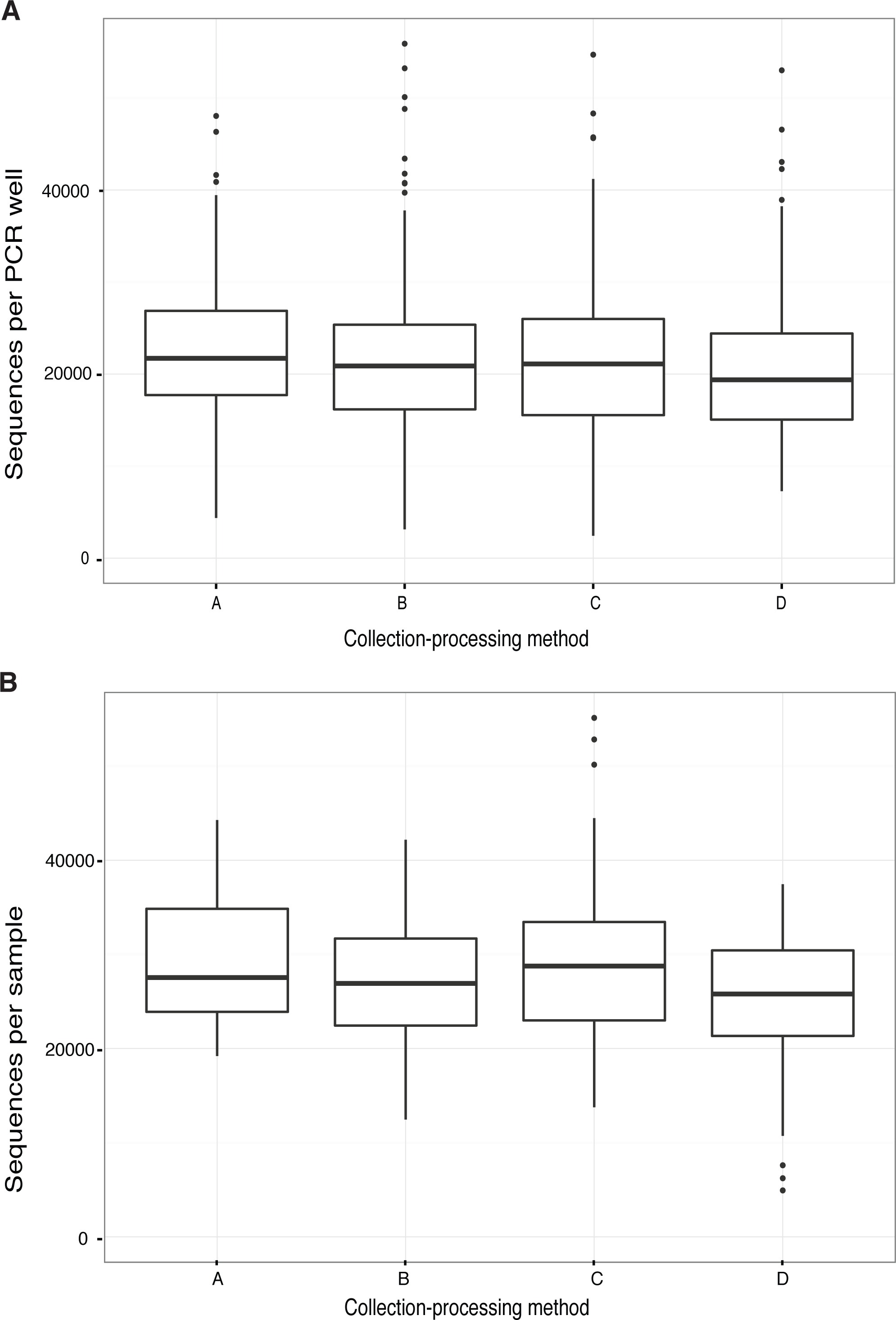
Library size by collection-processing method. (A) WEHI sequencing (B) BCM sequencing.

**Figure S6.**
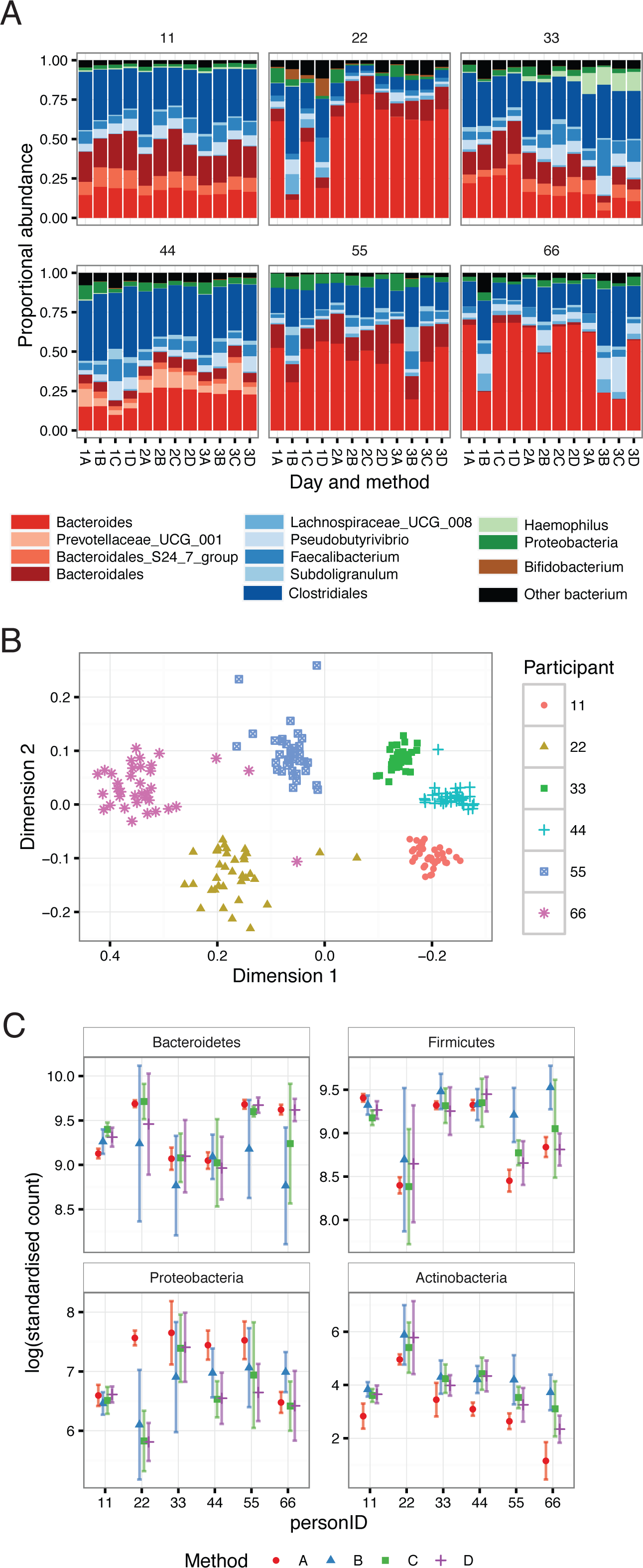
Overview of samples sequenced at Baylor College of Medicine and analysed with CMMR pipeline. (A) Stacked bar chart of dominant bacterial genera in each sample, equivalent to Figure 4A. Bars are colour-coded by phyla using the same colours as in Figure 2. Use of the Silva database for taxonomic assignment has introduced genus labels Prevotellaceae_UCG_001 and Lachnospiraceae_UCG_008. (B) Beta diversity using UniFrac distances between samples. Axes have been reflected to give approximately the same orientation of clusters as for Figure 2C. Directions in NMDS are arbitrary, so the positioning and rotation of clusters does not indicate a real change in the UniFrac distances. (C) Log of standardised counts (scaled by library size) of four phyla for the four methods for each individual, equivalent to Figure 3A. Points show mean and bars show standard deviation for each individual and collection-processing method (n=9).

## Abbreviations

ENDIA: Environmental determinants of islet autoimmunity
BCM: Baylor College of Medicine
WEHI: Walter and Eliza Hall Institute of Medical Research
16Sv4: hypervariable region of the 16S rRNA marker gene
OTU: Operational Taxonomic Unit
NMDS: Non-metric Multi-Dimensional Scaling
WW: WEHI bioinformatics analysis applied to DNA sequenced at WEHI
WB: WEHI bioinformatics analysis applied to DNA sequenced at BCM
BB: BCM bioinformatics analysis applied to DNA sequenced at BCM (supplementary material)

